# ELONGATED HYPOCOTYL 5 (HY5) and POPEYE (PYE) Regulate Intercellular Iron Transport in Plants

**DOI:** 10.1101/2024.05.06.592684

**Authors:** Samriti Mankotia, Abhishek Dubey, Pooja Jakhar, Santosh B. Satbhai

## Abstract

Plants maintain iron (Fe) homeostasis under varying environmental conditions by balancing processes such as Fe uptake, transport, and storage. In Arabidopsis, POPEYE (PYE), a basic helix-loop-helix (bHLH) transcription factor (TF), has been shown to play a crucial role in regulating this balance. In recent years, the mechanisms regulating Fe uptake have been well established but the upstream transcriptional regulators of Fe transport and storage are still poorly understood. In this study, we report that ELONGATED HYPOCOTYL5 (HY5), a basic leucine zipper (bZIP) TF which has recently been shown to play a crucial role in Fe homeostasis, interacts with PYE. Molecular, genetic and biochemical approaches revealed that PYE and HY5 have overlapping as well as some distinct roles in regulation of Fe deficiency response. We found that HY5 and PYE both act as a repressor of Fe transport genes such as *YSL3, FRD3 NPF5.9, YSL2, NAS4,* and *OPT3*. HY5 was found to directly bind on the promoter of these genes and regulate intercellular Fe transport. Further analysis revealed that HY5 and PYE directly interact at the same region on *PYE* and *NAS4* promoter. Overall, this study revealed that HY5 regulates Fe homeostasis by physically interacting with PYE as well as independently.

## INTRODUCTION

Iron (Fe) homeostasis is crucial for optimum growth and development of plants. Fe deficiency in plants affects photosynthesis, nutrient transport, immunity, and mitochondrial respiration which ultimately leads to defects in plant growth and development (Balk and Schaedler, 2014; Hänsch and Mendel, 2009). On the other hand, Fe excess is toxic for plants, and it leads to inhibition of root growth, leaf bronzing, necrosis, and biomass reduction (Zhang *et al*., 2018). Therefore, in order to maintain Fe homeostasis, plants have adapted various mechanisms which regulate Fe uptake, transport, storage, and assimilation.

In *Arabidopsis thaliana* (Arabidopsis) a reduction-based mechanism, also known as Strategy I, ensures optimum Fe uptake from the soil (Kobayashi and Nishizawa, 2012; Brumbarova *et al*., 2015). The Strategy I is mainly dependent on the activity of ATpase2 (AHA2), FERRIC REDUCTION OXIDASE2 (FRO2), and IRON-REGULATED TRANSPORTER1 (IRT1). The release of H^+^ ions in the rhizosphere by AHA2 decreases the pH of the rhizosphere, enhancing the solubility of insoluble Fe^3+^ present in the soil (Santi and Schmidt, 2009). The solubility of Fe^3+^ is also enhanced by the coumarins released into the rhizosphere by the PLEIOTROPIC DRUG RESISTANCE (PDR9) transporter (Fourcroy *et al*., 2014). FRO2 enzyme reduces this soluble Fe^3+^ into Fe^2+^ which is taken up by the roots with the help of IRT1 transporter (Ying Yi and Mary Lou Guerinot, 1996; Nigel j. Robinson *et al*., 1999; Eide *et al*., 1996; Vert *et al*., 2002). Once inside the root, Fe is translocated to the shoot, followed by redistribution to sink tissues, and excess Fe is stored in the vacuoles to avoid Fe toxicity. Fe is transported in chelated forms with citrate and nicotianamine (NA). Fe is transported from root to shoot through xylem in the form of Fe^3+^-citrate complex. FERRIC REDUCTASE DEFECTIVE 3 (FRD3) is involved in the transport of citrate in the xylem (Durrett *et al*., 2007). OLIGOPEPTIDE TRANSPORTER 3 (OPT3) mediates the transport of Fe from root to shoot, from xylem to phloem, and facilitates Fe distribution from source to sink tissues (Stacey *et al*., 2002; Stacey *et al*., 2008). In addition to OPT3, two NITRATE AND PEPTIDE TRANSPORTER FAMILY (NPF) transporters, NPF5.8 and NP5.9, have been shown to regulate long-distance Fe transport from root to shoot (Chen *et al*., 2021). YELLOW STRIPE-LIKE (YSL) proteins are responsible for the transport of Fe-NA complexes. YSL2 mediates the transport of Fe through vascular system (DiDonato *et al*., 2004). YSL1 and YSL3 are involved in transport of Fe to seeds and senescent leaves (Waters *et al*., 2006).

The Fe homeostasis is regulated at the transcriptional level by a cascade of basic leucine zipper (bHLH) transcription factors (TFs). The expression of major Fe uptake genes *IRT1, FRO2* and *AHA2* is induced under -Fe which is under the control of FIT and bHLH Ib (bHLH38, bHLH39, bHLH100, and bHLH101) TFs (Yuan *et al*., 2008; Wang *et al*., 2013; Liang *et al*., 2017). The expression of bHLH Ib TFs is also induced under -Fe by the bHLH IVc TFs (bHLH34, bHLH105/ILR3, bHLH104, and bHLH115) and bHLH121/URI (Li *et al*., 2016; Liang *et al*., 2017; Kim *et al*., 2019; Gao, Robe and Dubos, 2020). The bHLH IVc TFs are known to be regulated at the protein level by the activity of BRUTUS (BTS). BTS is a E3 ubiquitin ligase, and it degrades bHLH IVc TFs under Fe sufficient conditions to avoid its overload (Selote *et al*., 2015). Interestingly, IRON MAN PEPTIDES (IMAs) interact with BTS which stabilizes bHLH IVc TFs under Fe deficient conditions (Grillet *et al*., 2018; Li *et al*., 2021). POPEYE (PYE), a transcriptional factor belonging to bHLH IVb group is also upregulated under Fe deficiency (Long *et al*., 2010). PYE is known to act as transcriptional repressor to regulate Fe homeostasis in plants. The PYE-mediated transcriptional repression of genes under Fe deficiency is essential for maintaining Fe homeostasis. PYE is known to interact with ILR3 and directly negatively regulate the expression of *NAS4* (involved in iron transport)*, FER1, FER2, FER4* and *VTL2* (involved in iron storage) and *NEET* (iron assimilation) (Tissot *et al*., 2019). Even though earlier studies have revealed several upstream transcriptional regulators of Fe uptake, it is still unclear that which upstream transcriptional regulators are regulating the iron transport.. Previously, we have shown that ELONGATED HYPOCOTYL 5 (HY5), a bZIP TF act as both, transcriptional activator and repressor to regulate Fe deficiency responses. HY5 has been found to positively regulate the expression of Fe uptake genes which include *FRO2, IRT1,* and *FIT* and negatively regulate the expression of *BTS* and *PYE* (Mankotia *et al*., 2023). HY5 can function both as a transcriptional activator and repressor depending on its binding partner (Ang *et al*., 1998; Delker *et al*., 2014; Gangappa and Botto, 2016; Gangappa and Kumar, 2017; Burko *et al*., 2020). However, the binding partners of HY5 in regulating Fe homeostasis are not identified.

In this study, we performed yeast two-hybrid (Y2H) assay followed by bimolecular fluorescence complementation (BiFC) and found that HY5 interacts with PYE. We found that *pyehy5* double mutant was more sensitive to Fe deficiency as compared to both the single mutants. Transcriptional analysis together with chromatin immunoprecipitation (ChIP) assays revealed that HY5 also directly regulates the expression of genes involved in Fe transport including *YSL3, FRD3, NPF5.9, YSL2, NAS4,* and *OPT3.* Interestingly, HY5 and PYE were found to directly bind at the same region on *PYE* and *NAS4* promoter. Our findings support a model in which HY5 plays an essential role in regulation of Fe deficiency response by interacting with PYE and also independently possibly by interacting with other TFs. In addition, our findings shed light on the regulation of intercellular Fe transport by HY5.

## 2 MATERIALS AND METHODS

### 2.1 Plant materials and growth conditions

Columbia (Col-0) ecotype of Arabidopsis thaliana was used as WT. The mutant lines used in this study were *hy5* (SALK_056405) and *pye* (SALK_021217) in the Col-0 background. The *pHY5:HY5:YFP/hy5* line was provided by Roman Ulm. The *pPYE::PYE:GFP/pye* was provided by Terri Long (Long *et al*., 2010). The *hy5pye* double mutant was generated by crossing the *hy5* and *pye* single mutants. The mutants (single as well as double) were confirmed by performing genotyping PCR. The primers which are used for performing the genotyping PCR are listed in Supplemental Table 1. For seed sterilization, 5% sodium hypochlorite and 70% ethanol was used. After sterilization, seeds were kept in 4°C for 3 days. Seeds were sown on ½ Murashige and Skoog media (Caisson Labs, Smithfield, UT, USA) with Fe (+Fe) or without Fe (-Fe). The media pH was maintained 5.7 with KOH. For making media Fe deficient, 300 μM of [3-(2-pyridyl)-5,6-diphenyl-1,2,4-triazine sulfonate] Ferrozine (Sigma-Aldrich, Bangalore, India) was added in -Fe media. Seedlings were grown under long-day conditions, 16 h light and 8 h dark at 22 °C with a light intensity of 100– 110□μmol□cm^−2^□sec^−1^ and 50% humidity. The soilrite, perlite and compost (3:1:1) mixture was used as soil.

### 2.2 Phenotypic studies

For phenotypic study, the seeds were grown on ½ MS media containing Fe (+Fe) and on ½ MS media without Fe (-Fe). In the growth chamber, the plates were placed vertically and after 10-days, plates were scanned using the Epson Perfection V600 with 1200 dpi resolution. The root length quantification was performed using ImageJ 1.52a software (National Institutes of Health).

### 2.3 Chlorophyll content determination

For the measurement of chlorophyll content seedling were grown on +Fe and -Fe for 10 days. The chlorophyll was extracted from the leaf tissue of five-six seedlings using 1ml 80% acetone and incubated in dark for 24 h. The chlorophyll content was measured using the formula: (mg/g)□=□(20.3□×□*A*_645_□±□8.04□×□*A*_663_)□×□*V*/*W*□×□10^3^ (Aono *et al*., 1993).

### 2.4 Fe staining

For Fe staining, seedlings grown on +Fe for 5 days were used, and vacuum infiltrated with a solution of 1% (v/v) HCl and 1% (w/v) K-ferrocyanide for 5 min. The seedlings were washed using water for 4 to 5 times and imaged using NIKON ECLIPSE Ni U microscope.

The diaminobenzidine (DAB) intensification was done after Perls staining and for this, seedlings were incubated in methanol containing 10mM Na-Azide and 0.3% (v/v) H_2_O_2_ for 1 hour. After 1 hour incubation, the seedlings were washed using 100mM Na phosphate buffer (pH 7.4) and again incubated in same buffer which contains 0.025% (w/v) DAB and 0.005% (v/v) H_2_O_2_. The reaction was stopped by washing the seedlings with water. The images of stained seedlings were captured using NIKON ECLIPSE Ni U microscope.

### 2.5 Targeted Yeast Two-Hybrid (Y2H)

For performing Y2H, the AD (pDEST/GADKT7) and BD (pDEST/pGBKT7) constructs of the selected genes were generated by gateway cloning. The constructs were co-transformed into PJ697a, followed by plating on selection plate −2 (-Leu, -Trp). The colonies were obtained after 2 to 3 days, which were dissolved in 100μl of autoclaved millipore water. O.D_.600_ was adjusted to 2.0 in a 96-well plate and spotting was done on −4 (-Leu, -Trp, -His, - Ade) selection plates. The plates were scanned after 6 to 7 days to check growth and identify interacting colonies.

### 2.6 Cloning for BiFC

The CDS of gene of interest was first cloned in pENTR/D/TOPO vector. The pENTR/D/TOPO construct of the gene of interest was used to set up LR reaction with the gateway compatible destination vector pSITE BiFC cEYFP or pSITE BiFC nEYFP. The translational fusion was confirmed by performing restriction digestion and sequencing.

### 2.7 Bimolecular Fluorescence Complementation Assay

The *Nicotiana benthamiana* was grown in 16 hrs light and 8 hrs dark conditions for the bifluorescent complementation assay. The leaves of a 2-week-old plant were used for Agrobacterium-infiltration. After 3 days of infiltration, the leaves were screened with the help of a confocal microscope. The vectors containing the N- and C-terminal fragments of YFP respectively were utilized for the generation of N-terminal and C-terminal fusion of PYE and HY5. The constructs of nYFP and cYFP-HY5, nYFP-HY5 and cYFP-HY5, nYFP-PYE and cYFP or of nYFP-PYE and cYFP-HY5 were transformed into Agrobacterium strain GV3101.

### 2.8 Agrobacterium-mediated infiltration

The various constructs in the Agrobacterium were separately inoculated in 5ml LB containing rifampicin, gentamycin, and spectinomycin. The Agrobacterium having p19 RNAi suppressor gene construct was also inoculated similarly in 5ml LB containing rifampicin, gentamycin, and kanamycin. All the cultures were kept in shaker at 30 ℃ for growth. After 24 hours, the cultures were centrifuged at 4000rpm for 15 minutes. After centrifugation, the supernatant was discarded, and the pellets were washed twice in 2ml of infiltration buffer composed of 10mM Mgcl2, 10mM MES/KOH (pH 5.6), 150µM Acetosyringone. The pellet was finally resuspended in infiltration buffer and OD600 was adjusted in the range of 0.8 to 1.0. The resuspended bacterial cells were finally mixed in a 1:1:1 ratio in a 2ml MCT and kept for 1 hour incubation at 30℃. The resulting mixture was injected in the abaxial surface of a leaf of *Nicotiana benthamiana.* The plants were kept at 22℃ and leaves were screened after 2 to 3 days using a confocal microscope (Leica SP8 upright).

### 2.9 Confocal microscopy

For confocal microscopy, the leaves were stained with Propidium Iodide (100 µg/ml) in dH20. The leaves were then scanned using Leica SP8 upright laser scanning confocal microscope. Argon laser (488-nm) was used to excite the YFP and Propidium Iodide was excited using 561-nm laser. The emission spectra was collected at 500-530 nm and 600-650 nm, respectively.

### 2.10 RNA isolation and qRT-PCR

The total RNA was isolated from the roots of seedlings which were grown for 6 days on ½ MS and transferred to +Fe or -Fe for 3 days using the Qiagen Plant RNeasy kit following the manufacturer’s protocol. The isolated RNA was treated with DNase I for the removal of genomic DNA. The cDNA was synthesized from 2µg RNA using the RevertAidTM First Strand cDNA synthesis kit (Thermo). The qPCR was set up using the LightCycler 480 II (Roche) and TB GreenTM Premix Ex. The relative expression levels were measured using the comparative threshold cycle method (ΔΔCT) and for the reference gene, β-tubulin was used. The primers used are listed in Supplemental Table 1. The 3µg of isolated RNA was sent for RNA Sequencing.

### 2.11 RNA-Sequencing data analysis

RNA-Seq libraries were prepared using NebNext Ultra II RNA library prep kit for Illumina(E7770L) according to manufacturer’s protocols and quality and quantity was assessed using Agilent 4150 TapeStation system and Qubit 4 Fluorometer. High quality Total RNA-Seq libraries were sequenced (2×150 bp) on NovaSeq 6000 V1.5. Bulk RNA sequencing data was analyzed as per the standard practices. Briefly, QC was performed on all fastq files using FASTQC following which 5’ sequences and adapter sequences were removed using trimmomatic (Bolger *et al*., 2014). The output fastq files were mapped to Arabidopsis thaliana genome (TAIR10) using HISAT2 (Zhang *et al*., 2021). Subsequently, the counts file was generated using featureCounts (Liao *et al*., 2014). All downstream analysis was performed in a Python environment. Counts files for all the samples were imported as a dataframe and subjected to basic filtering in which genes having counts less than 10 were removed. Thereafter, FPKM was calculated for all the genes and a second filtering step was applied to remove genes having FPKM < 0.5. Seaborn and matplotlib was used for plotting. Counts file was used to perform DEGs analysis using pyDESEQ2. A p-value cutoff of 0.05 and Fold Change cutoff of 1.5 was applied to identify significant DEGs.

### 2.12 ChIP assay

The ChIP experiment was conducted according to a previously described method (Gendrel *et al*., 2005). The *pHY5:HY5:YFP/hy5* and *35S:eGFP* was in the Ler background. The *pPYE:PYE:GFP/pye* and *35S:eGFP* was in the Columbia background. The seedlings were germinated and grown on ½ MS media for 10 days. The fixation was done using 1% formaldehyde. The nuclei isolation was done followed by sonication using Qsonica 800R ultrasound sonicator with around 30 cycles of 70% pulse amplitude for 15 sec followed by 45 sec of pulse off time. The protein-G magnetic beads (10004D; Dynabeads) and anti-GFP antibody (a290; Abcam) were used to pull down protein-DNA complexes. IgG was used as a negative control. The beads were washed and the immunocomplexes bound to the beads were eluted and reverse crosslinked. The precipitated chromatin was used to set up qPCR. For calculating Fold enrichment, normalization was done against the negative control, i.e., IgG. The primers used are mentioned in Supplemental Table 1.

### 2.13 Statistical analysis

Data was expressed as means of ± SEM. P-values were determined using Student’s t-test or one-way ANOVA followed by *post hoc* Tukey HSD test. The P-value of ≤ 0.05 was considered as statistically significant.

## 3 RESULTS

### 3.1 HY5 Physically interacts with PYE

Arabidopsis HY5 has been known to act both as a transcriptional activator and repressor depending on its binding partner (Ang *et al*., 1998; Burko *et al*., 2020; Delker *et al*., 2014; Gangappa and Kumar, 2017; Gangappa and Botto, 2016; Nawkar *et al*., 2017; Ruckle *et al*., 2007; Xu *et al*., 2016; Yadukrishnan *et al*., 2020; Zhang *et al*., 2017). In BioGRID database 48 interactors of HY5, including PYE and some members of subgroup Ib transcription factors, bHLH39 and bHLH101 are reported (Stark *et al*., 2006). We therefore performed Y2H of HY5 with the major bHLH TFs involved in regulating the iron deficiency response to identify potential interacting partners of HY5. Y2H assays were performed using HY5 fused to the binding domain of GAL4 as bait and bHLH TFs (bHLH34, bHLH105/ILR3, bHLH104, bHLH115, bHLH38, bHLH39, bHLH100, bHLH101, PYE, FIT, BTS) fused to the activation domain of GAL4 as prey. HY5 fused to the activation domain of GAL4 was used as a positive control as HY5 is known to interact with itself to form homodimers. We found that HY5 interacts with some of the bHLH TFs in one or two Y2H assays but only PYE was found to be interacting with HY5 consistently in all the three independent assays (Figure 1a). To further confirm our results, the interaction between HY5 and PYE was further analyzed using bimolecular fluorescence complementation (BiFC) assays. For the BiFC assays, PYE was fused to the N-terminal part of YFP (nYFP-PYE) and HY5 was fused with C-terminal part of YFP (HY5-cYFP). HY5 was also fused with N-terminal part of YFP (nYFP-HY5) which was used as a positive control. We observed strong signal when HY5-cYFP was assayed with nYFP-PYE and nYFP-HY5, whereas no signal was observed when HY5-cYFP was assayed with nYFP and nYFP-PYE was assayed with cYFP (Figure 1b). Altogether, these results demonstrate that HY5 interacts with PYE and can form heterodimer with PYE.

**Figure 1.**
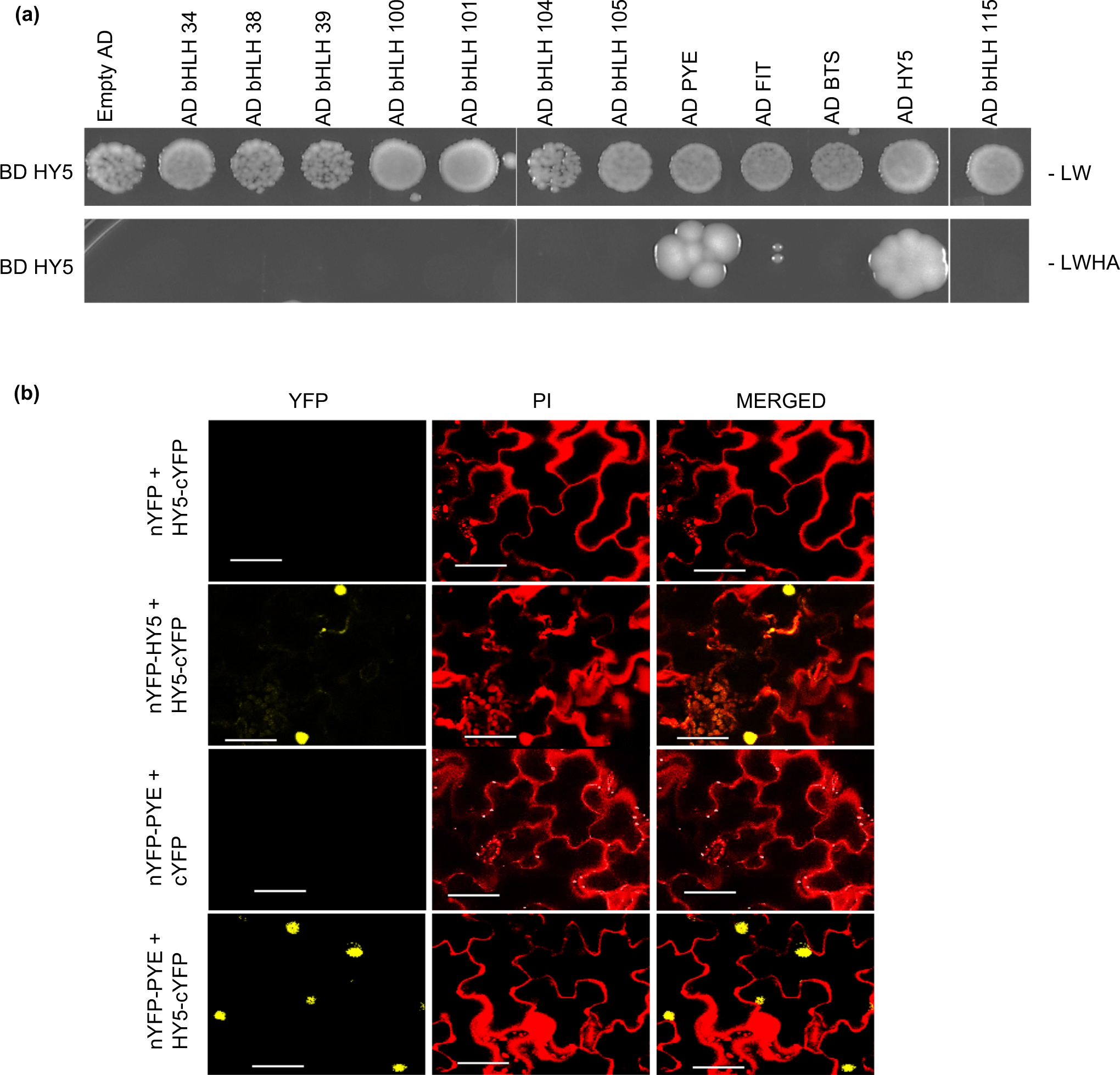
HY5 and PYE play additive roles in regulating iron deficiency response. **(a)** Y2H assays. bHLH34, bHLH38, bHLH39, bHLH100, bHLH101, bHLH104, bHLH105, PYE, FIT, BTS and bHLH115 fused with the GAL4 activation domain (AD) and HY5 with the GAL4 binding domain (BD) were cotransformed in the yeast strain. The different yeast strains were plated on non-selective media (-LW) and also on selective media (-LWHA). AD alone was used as negative control. The growing colonies on the selective media represent positive Y2H interacting colonies and they were identified after 7 days of growth. H, histidine; L, leucine; W, tryptophan, A, adenine. **(b)** BiFC assays. PYE was fused with N-terminal part of YFP(nYFP) and HY5 was fused with both N-terminal and C-terminal part of YFP (cYFP) prior to infiltration into Nicotiana benthamiana leaves and analysis was done using confocal microscopy. nYFP and cYFP alone were used as a negative control. Bar=100µM.

### 3.2 Genetic interaction exists between HY5 and PYE

Considering the finding that HY5 physically interacts with PYE; we investigated their genetic interaction. For this, we generated *hy5pye* double mutant by crossing the corresponding single mutants and compared its phenotype with the wild type (WT) as well as both the *hy5* and *pye* single mutants. When grown on -Fe, both the *hy5* and *pye* single mutants were more sensitive to -Fe as compared to the WT which is in accordance with the previously published reports (Long *et al*., 2010; Mankotia *et al*., 2023) (Figure 2a). Furthermore, we observed that the *hy5pye* double mutant was highly sensitive under -Fe as compared to both the single mutants and had significantly shorter roots when grown on -Fe as compared to +Fe conditions (Figure 2b). The percentage reduction in the chlorophyll content of *hy5pye* double mutant grown on -Fe as compared to those grown on +Fe was also found to be higher as compared to both the single mutants (Figure 2c). Next, we checked Fe accumulation in the roots of *hy5pye* double mutant and compared it with the corresponding single mutants and WT. We found that the Fe content is less in the *hy5* mutant, more in the *pye* mutant as compared to the WT which is in accordance with previously published reports and in the double mutant Fe content was less as compared to the WT and *pye* single mutant (Long *et al*., 2010) (Figure S1a,b). Both *hy5* and *pye* mutant showed similar phenotypes under -Fe and the double mutant was more sensitive as compared to both the single mutants. These results suggest that HY5 and PYE play additive roles in Fe deficiency response. We also hypothesized that these two genes might have synergistic as well as independent roles in regulation of Fe deficiency response.

**Figure 2.**
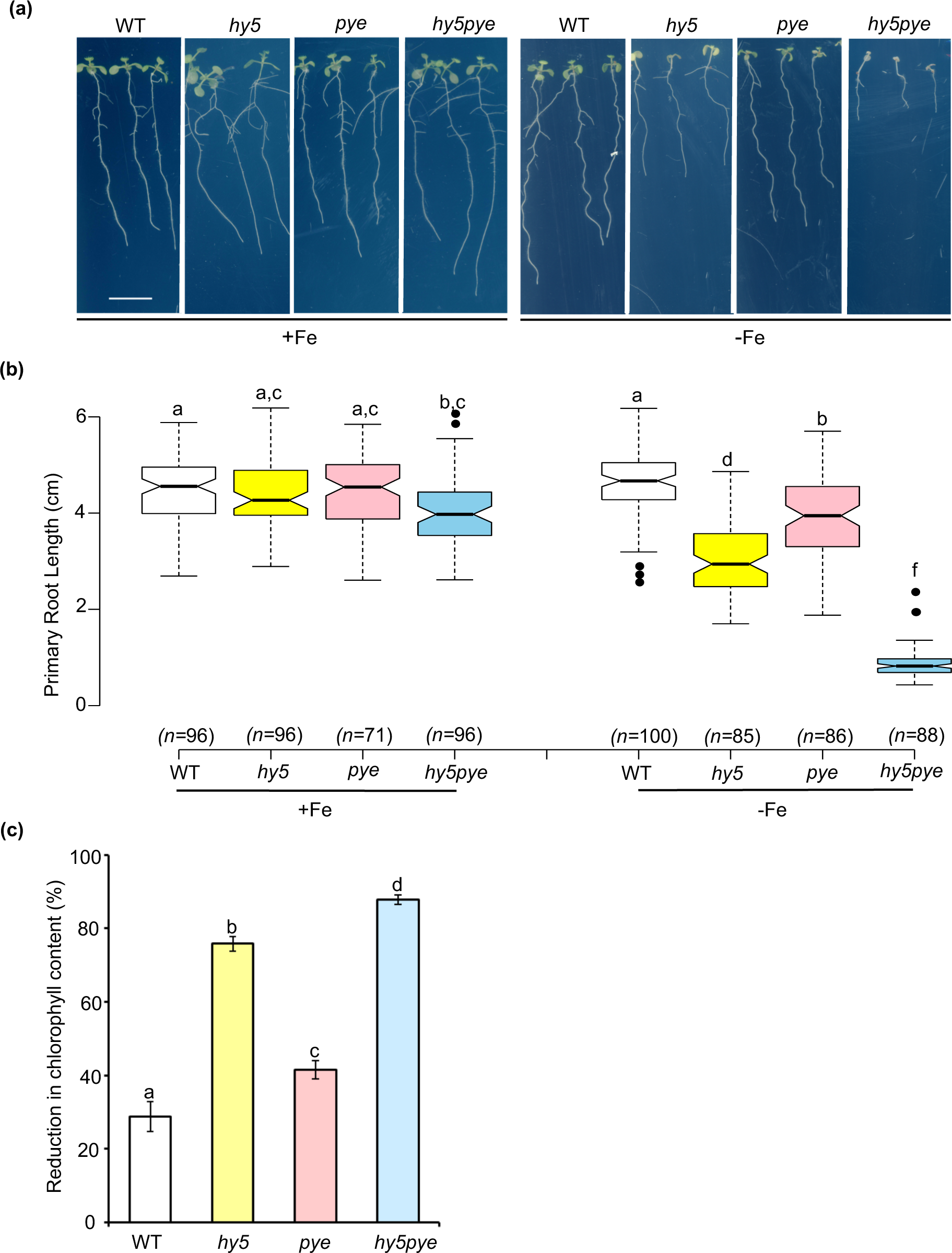
*HY5* and *PYE* play additive roles in regulating iron deficiency response. **(a)** Phenotypes of the *Arabidopsis* wild type (WT), *hy5*, *pye* and *hy5pye* grown for 10 days on Fe-sufficient and Fe-deficient medium. Scale bars, 0.5cm. **(b)** Boxplot of **r**oot length of the wild type (WT), *hy5*, *pye* and *hy5pye* grown for 10 days on Fe-sufficient and Fe-deficient medium. Means within each condition with the same letter are not significantly different according to one-way ANOVA followed by post hoc Tukey test, P < 0.05. **(c)** Percentage reduction in Chlorophyll content of the wild type (WT), *hy5*, *pye* and *hy5pye* grown grown on –Fe as compared to those grown on +Fe. Data shown is an average of three independent experiments. Each experiment consists of 4 biological replicates and each replicate consists of a pool of around six seedlings. Error bars represent ±SEM. Means within each condition with the same letter are not significantly different according to one-way ANOVA followed by post hoc Tukey test, P < 0.05.

### 3.3 Genome-wide expression analysis showed mis-regulation of Fe deficiency-responsive genes in the *hy5, pye* and *pyehy5* mutant

Since, HY5 interacts with PYE both physically and genetically. We next wanted to find out the genes regulated by HY5 and PYE together as well as independently. For this, we analyzed the transcriptomic changes in the roots of *pye, hy5, hy5pye* and WT that were grown on +Fe conditions for 6 days and exposed to +Fe or -Fe (+300μM Ferrozine) conditions for 3 days before harvesting. We compared the differentially expressed genes in the *pye, hy5,* and *hy5pye* with the WT (-Fe vs +Fe). We used 1.5-fold difference in gene expression as a cutoff. We found that 1,583 were differentially expressed genes (DEGs) between the *hy5* T (-Fe) vs WT T (-Fe), 2,791 genes were differentially expressed in the *pye* T (-Fe) vs WT T (-Fe), and 5,674 genes were differentially expressed in the *hy5pye* double mutant(-Fe) vs WT T (-Fe) (Figure 3a). Among these, 555 genes were the common DEGs in the *pye, hy5,* and *hy5pye* mutant (Figure 3a). Out of common DEGs, 254 genes were commonly upregulated in all, and 195 genes were commonly downregulated in all (Figure 3b, c). The genes involved in Fe uptake (*FIT, GRF11, IRT2, F6’H1* and *BGLU42)* were found to be commonly downregulated in the *hy5*, *pye* single as well as *hy5pye* double mutant (Figure 3b, d). The genes involved in Fe transport (*NPF5.9, FRD3* and *YSL3)* were found to be commonly upregulated in all (Figure 3c, d). To further explore the RNA-seq data, we performed Gene Ontology (GO) analysis using panther for the commonly differentially upregulated as well as downregulated genes in the *hy5* (T), *pye* (T) and *hy5pye* (T) as compared to the WT (T). This was conducted using genes that were upregulated or downregulated with more than 1.5-fold change. Major GO clusters for all commonly differentially expressed genes are shown in Figure 4. The response to iron ion starvation, cellular response to external stimulus, and response to light intensity were among the enriched downregulated GO terms. On the other hand, the nitrate metabolic process, glutathione metabolic process and nitrate assimilation were among the enriched upregulated GO terms. (Figure 4).

**Figure 3.**
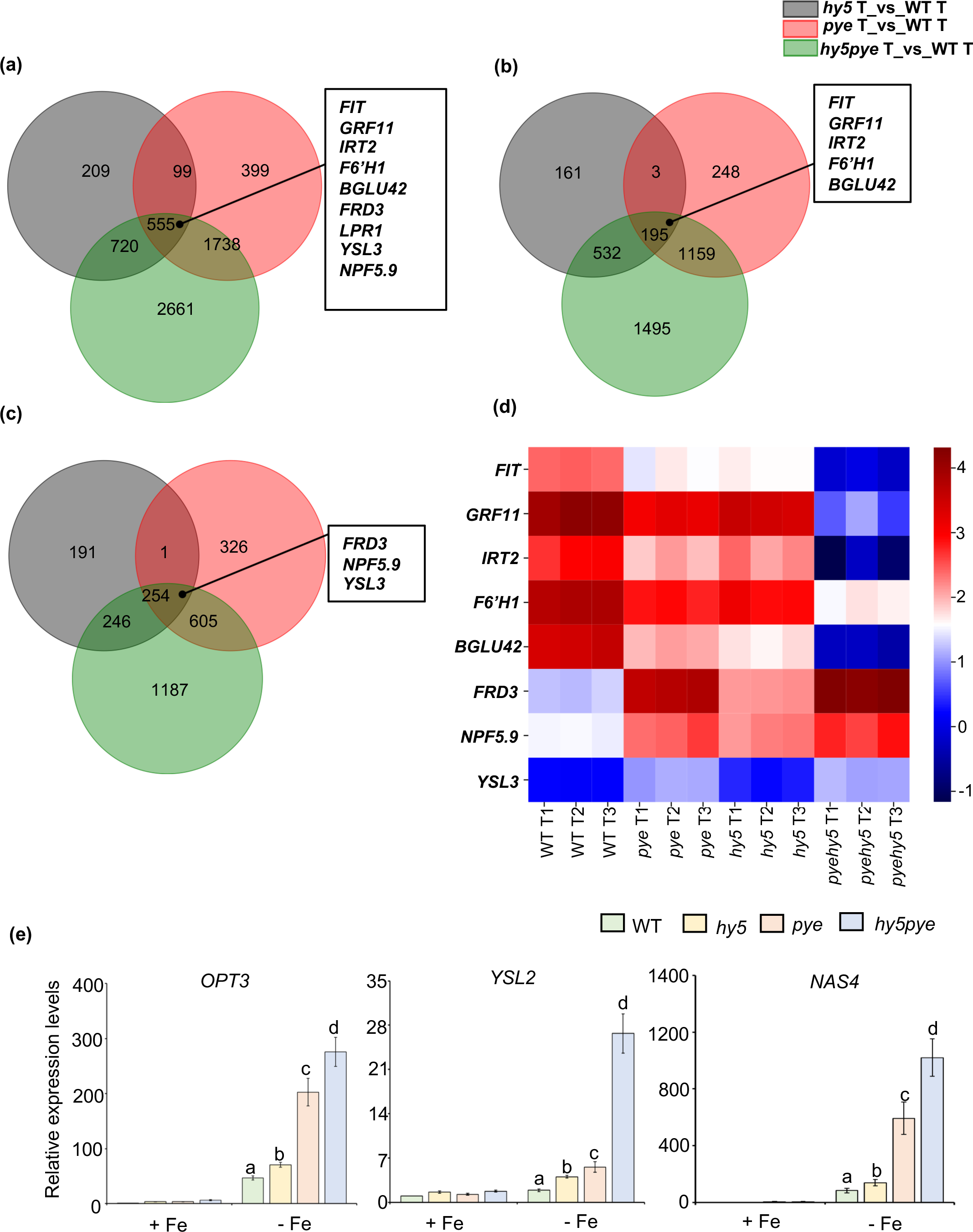
Fe deficiency-responsive genes are misregulated in the *pye, hy5* and *hy5pye* double mutant. **(a)** Venn diagram showing overlap of differentially expressed genes (DEGs; fold change; |FC| > 1.5) in the *hy5*, *pye* and *hy5pye* (T) versus WT (T). **(b)** Venn diagram showing overlap of commonly downregulated genes in the *hy5*, *pye* and *hy5pye* (T) versus WT (T). **(c)** Venn diagram showing overlap of commonly upregulated genes in the *hy5*, *pye* and *hy5* (T) versus WT (T). **(d)** Heat map of selected Fe deficiency responsive genes. The color bar on the right side demonstrates the log2 (FoldChange) of a given gene in response to iron deficiency for a given genotype. **(e)** Expression levels of *OPT3, YSL2*, and *NAS4.* Relative expression was determined by qRT-PCR in WT, *hy5, pye* and *hy5pye* double mutant grown on +Fe for days, and then transferred to both +Fe and –Fe (+300µM Fz) for 3 days. Data shown is an average of three biological replicates (n=2 technical replicates). Each biological replicate comprises of pooled RNA extracts from roots of 90 seedlings. Error bars represent ±SEM. *Significant difference by Student’s *t*-test (*P*□≤□0.05).

**Figure 4.**
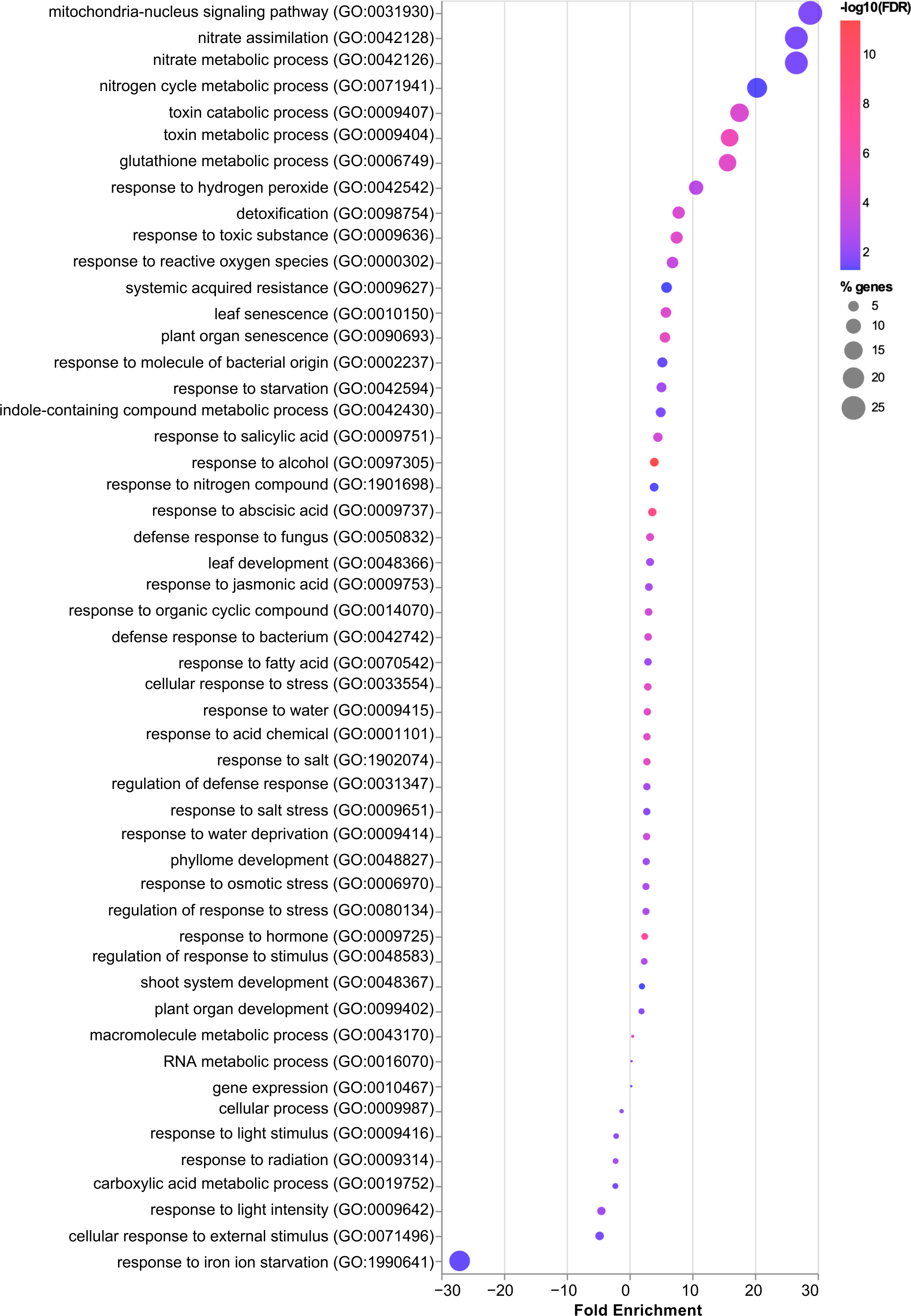
GO analysis of the differentially expressed genes. **(a)** Gene ontology analysis of the genes commonly differentially expressed in the *hy5* (T), *pye* (T) and *hy5pye* (T) as compared to the WT (T).

PYE is known to act as a transcriptional repressor and HY5 is known to act both as a transcriptional activator and repressor to regulate the expression of genes involved in Fe deficiency response (Long *et al*., 2010; Burko *et al*., 2020). So, we next compared the expression of genes which are known to be regulated by either PYE or HY5 in the *pye*, *hy5* single as well as *pyehy5* double mutant to find out whether HY5 has any impact on the genes regulated by PYE and vice-versa. We found that the genes involved in Fe uptake including *FRO2* and *IRT1* which are known to be positively regulated by HY5 were less induced in the *hy5* single mutant (T) which is in accordance with a previous study and they were not induced similar to the WT (T) levels in the *pye* mutant (T) (Figure S2) (Long *et al*., 2010; Mankotia *et al*., 2023). In the *hy5pye* double mutant (T) also, we observed that these genes were even much less induced ((Figure S2). The genes such as *NAS4, ZIF1*, *NEET*, and *FRO3* were found to be more induced and the genes such as *FER1, FER3,* and *FER4* were found to be less repressed in the *pye* single mutant as compared to the WT under Fe deficiency which is in accordance with previous studies (Long *et al*., 2010; Tissot *et al*., 2019). In the *hy5* mutant, the expression of *NAS4,* and *ZIF1* appears to be more induced as compared to the WT but not highly induced like the *pye* and *hy5pye* double mutant (Figure S2). *NAS4* is known to be involved in Fe transport, *FER1, FER3,* and *FER4* are known to be involved in Fe storage, *FRO3* is proposed to be involved in mitochondrial iron homeostasis, and *ZIF1* is known to be involved in vacuolar sequestration of Zinc (Klatte *et al*., 2009; Ravet *et al*., 2009; Petit *et al*., 2001; Haydon and Cobbett, 2007; Mukherjee *et al*., 2006; Heazlewood *et al*., 2004). The expression of *FER3, NEET*, *FER4*, and *FRO3* was not found to be induced more or less repressed in the *hy5* mutant as compared to the WT unlike *pye* mutant (Figure S2). As we have found that most of the Fe transport related genes were strongly induced in the *hy5, pye* and *hy5pye* mutant. Therefore, we also checked the expression of *OPT3* and *YSL2* as these genes are also known to play an important role in Fe transport and we found that these are also more induced in the *hy5*, *pye* and *hy5pye* double mutant. We next performed qRT-PCR to confirm the RNA-seq results as *OPT3, YSL2,* and *NAS4* expression appears to be only slightly more induced in the *hy5* mutant as compared to the WT. We found that similar to the results of RNA-seq, these genes were more upregulated in the *hy5* mutant as compared to the WT but the induction was not high as in the *pye* and the *hy5pye* double mutant (Figure 3e, S2). In addition to these genes, *BTS* which is a E3 ubiquitin ligase, and a negative regulator of Fe uptake genes was also found to be more upregulated in the *hy5* and *pye* mutant which is in accordance with the previously published reports (Figure S2). Altogether, these results demonstrate that the expression of several key genes involved in Fe uptake and transport was affected in both *pye* and *hy5* mutants in a similar manner but the expression of some of the genes involved in Fe uptake such as *FRO2*, and *IRT1* was only affected in the *hy5* mutant and some of the genes related to Fe storage and assimilation such as *FER1, FER3, FER4,* and *NEET* was only affected specifically in the *pye* mutant. These results further provide evidence that HY5 and PYE have some similar as well as some unique roles in the regulation of Fe deficiency response.

### 3.4 HY5 directly regulates the expression of genes involved in Fe transport

Next, we wanted to check whether HY5 directly binds to the promoter of *NPF5.9, FRD3, YSL3, YSL2* and *OPT3* to regulate their expression. For this, we performed in silico analysis of the promoter sequence of *NPF5.9, FRD3, YSL3, YSL2* and *OPT3*. We found HY5 binding motifs on the promoter of these genes (Figure 5a). The *YSL3* gene has one E-box which is 189 bp upstream (region I), one CA-Hybrid which is 870 bp upstream (region II) to ATG. The *FRD3* promoter has one T/G-box which is 116 bp upstream (region I), one ACE-element which is 1713 bp (region II) upstream to ATG. The *NPF5.9* promoter has one G-box which is 1529 bp upstream (region I) and one E-box which is 1961 bp (region II) upstream to ATG. The *OPT3* promoter has one G-box which is 145 bp upstream (region I), one G-box which is 344 bp upstream (region II), one T/G-box which is 444 bp upstream (region III), one G-box which is 575 bp ustream, one T/G-box which is 592 bp upstream, one G-box which is 622 bp upstream (region IV) and one CG-Hybrid which is 1056 bp (region V) upstream to ATG. The *YSL2* gene has one A-box which is 904 bp upstream to ATG (Figure 5a). Chromatin immunoprecipitation (ChIP)-qPCR was performed using anti-GFP antibody to check whether HY5 directly binds to the promoter of these genes to regulate the expression or not. The *Pro:HY5:HY5:YFP/hy5* and *35S:eGFP* were used for the ChIP experiments. The ChIP-qPCR results revealed that HY5 directly binds to the promoter of *FRD3, YSL3, NPF5.9* at region II (ACE-element, CA-Hybrid box, and E-box respectively), *YSL2* at region I (A-box) and *OPT3* at region IV (G-box, T/G box) (Figure 5b).

**Figure 5.**
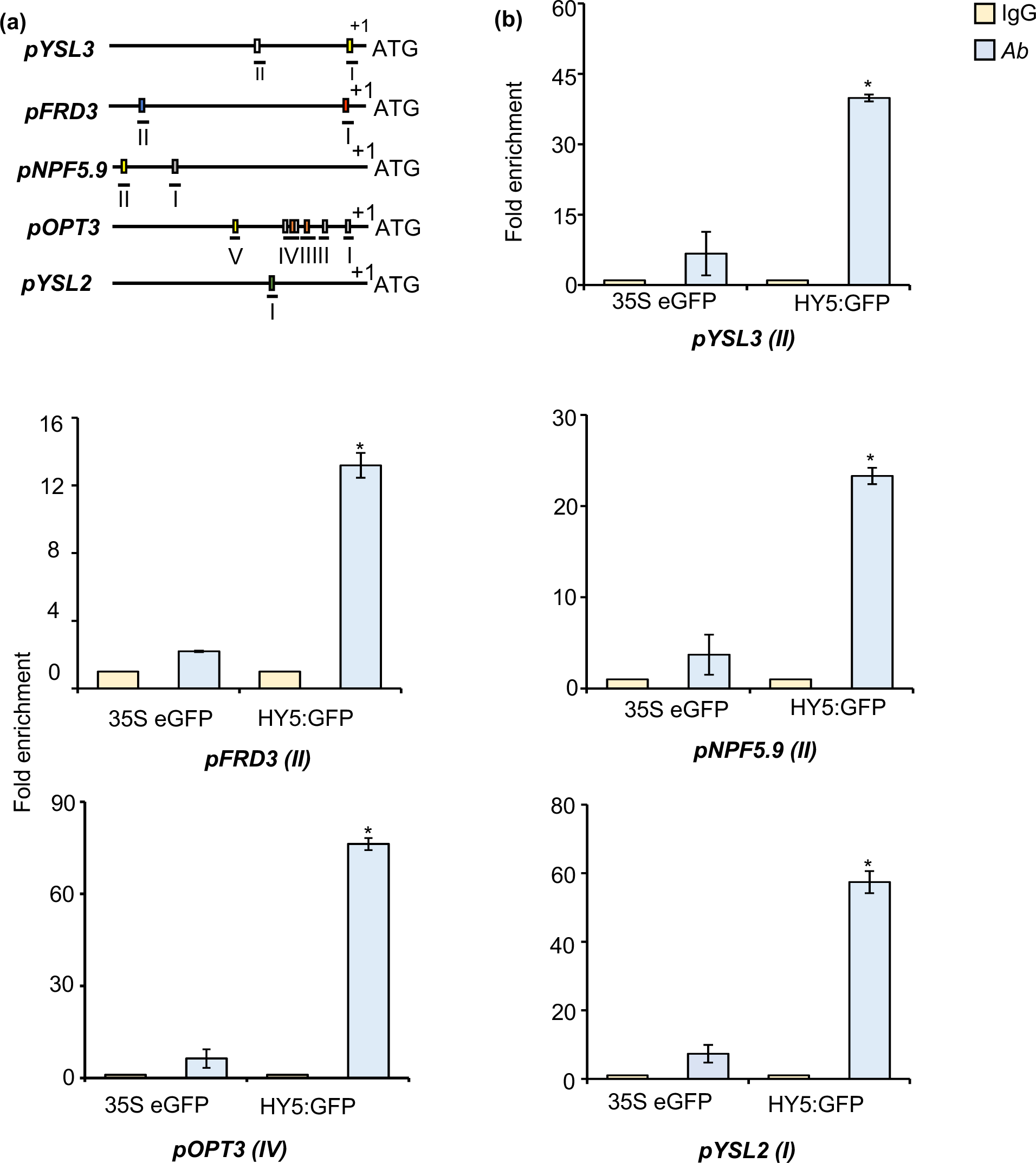
HY5 directly binds on *YSL3, FRD3, NPF5.9, OPT3 and YSL2* promoter. **(a)** Schematic diagram of the promoter of the *YSL3, FRD3, NPF5.9, OPT3* and *YSL2.* White box represent CA-Hybrid boxes, grey boxes represent G-boxes, orange boxes represent T/G-boxes, blue boxes represent ACE-element, green boxes represent A-boxes and yellow boxes represent E-boxes. Lines under the boxes represent sequences detected by ChIP qPCR. **(b)** ChIP-qPCR showing relative enrichment of the *YSL3, FRD3, NPF5.9, OPT3* and *YSL2* regulatory regions bound by HY5. The ChIP assay was performed using *pHY5::HY5:YFP/hy5* and *35S:eGFP* seedlings grown on +Fe media for 10 days. Error bars represent ±SEM. *Significant difference by Student’s t test (P≤0.05).

### 3.5 HY5 and PYE interact with the same promoter loci on *PYE* and *NAS4* promoter

We found that HY5 directly regulates *OPT3, YSL2, YSL3, FRD3* and *NPF5.9* and the expression of these genes was also found to be more induced in the *pye* mutant. PYE is known to bind G-box and E-box (De Masi *et al*., 2011). So, we hypothesized that HY5 and PYE might together interact *in planta* with the same promoter loci on *OPT3* and *NPF5.9* to regulate the expression of these genes. For this, ChIP experiments were performed on the transgenic line *Pro:PYE:PYE:GFP/pye* using the anti-GFP antibody. The results revealed that PYE does not interact with the same promoter loci as HY5 does on *OPT3* and *NPF5.9* (Figure 6a). Therefore, PYE might regulate the expression of these genes indirectly.

**Figure 6.**
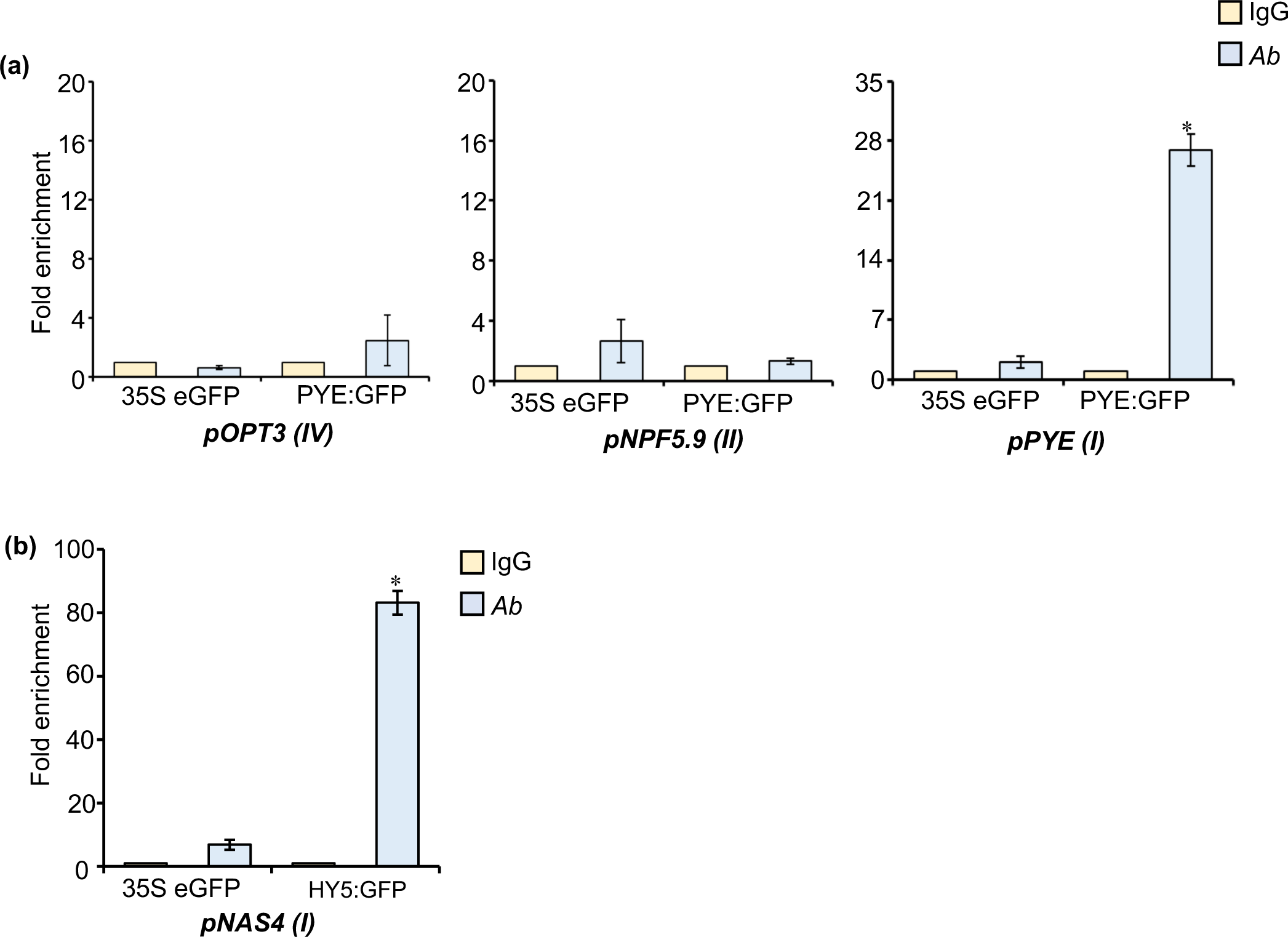
PYE and HY5 directly interact at same region on PYE and NAS4 promoter. **(a)** ChIP-qPCR analysis of the binding of the PYE to the promoter regions of the *OPT3, NPF5.9* and *PYE* that are targeted by HY5. The ChIP assay was performed using *pPYE::PYE:GFP/pye* and 35S:eGFP seedlings grown on +Fe media for 10 days. Error bars represent ±SEM. *Significant difference by Student’s t test (P≤0.05). **(b)** ChIP-qPCR analysis of the binding of the HY5 to the promoter region of the *NAS4* which is targeted by PYE. The ChIP assay was performed using *pHY5::HY5:YFP/hy5* and 35S:eGFP seedlings grown on +Fe media for 10 days.

In our previous study, we have found that HY5 binds to the region I on *PYE* promoter and negatively regulates its expression. It is known that ILR3 and PYE also interacts with the promoter of *PYE* (Tissot *et al*., 2019). So, next we wanted to confirm whether PYE also binds to same region I to which HY5 binds. For this, we performed ChIP experiment and we found that PYE also interacts at the same locus as HY5 does on the *PYE* promoter (Figure 6a).

As we have found that *NAS4* which is a known direct target of PYE is more induced in the *hy5* mutant as compared to the WT (Tissot *et al*., 2019) (Figure 3e). Next, we wanted to check whether HY5 also directly binds to the *NAS4* promoter region on which PYE binds. To confirm this, we performed ChIP experiment using *Pro:HY5:HY5:YFP/hy5* and *35S:eGFP*. We found that HY5 also interacts at the same region on *NAS4* promoter to which PYE binds (Figure 6b).

## 4 DISCUSSION

Although the molecular mechanisms regulating Fe uptake from the soil are relatively well studied, very little is known about how plants regulate cellular Fe homeostasis and intercellular Fe transport. An extensive regulatory network comprising of numerous bHLH TFs is essential for the regulation of Fe homeostasis in plants. Fe uptake, transport, storage, and assimilation must be tightly regulated in order to meet Fe requirements in various tissues and organs. PYE, a bHLH TF belonging to subgroup IVb is known to act as transcriptional repressor to regulate the expression of genes involved in internal mobilization of Fe, transport of Fe from root to shoot, and Fe storage (Long *et al*., 2010; Tissot *et al*., 2019). Recently, HY5 belonging to bZIP TF family has been found to regulate several genes involved in Fe uptake (Mankotia *et al*., 2023). However, it is still unclear whether HY5 also regulates other Fe transport or storage related genes or not.

In this study, we carried out Y2H assay followed by BiFC to identify the interacting partners of HY5. The results revealed that HY5 interacts with PYE (Figure 1a, b). The phenotyping experiments showed that both the *hy5* and *pye* mutants were sensitive to Fe deficiency but *hy5pye* double mutant was more sensitive to Fe deficiency as compared to both the single mutants under -Fe (Figure 2). Fe content analysis by both Perl’s and Perls/Dab staining revealed that Fe content was more in the *pye* mutant and less in the *hy5* mutant as compared to the WT (Figure S1). Transcriptional analysis revealed that the transcriptional response to Fe deficiency was altered in the *hy5, pye,* and *hy5pye* double mutant (Figure 3, S2). Some of the Fe uptake genes were less induced in the *hy5* as well as *pye* single mutants and they were even more less induced in the *hy5pye* double mutant under -Fe conditions as compared to the WT (Figure 3, S2). Inversely, the Fe transport related genes were found to be more induced in the *pye, hy5,* and *hy5pye* double mutant under -Fe conditions as compared to the WT (Figure 3, S2). The expression of some genes such as *FER1, FER3, FER4, NEET, FRO2,* and *IRT1* was affected only in the *pye* or *hy5* mutant and not in both the mutants (Figure S2). The *hy5* mutant has less iron content as compared to the WT and this might be the reason that the expression of Fe storage and Fe assimilation related genes such as *FER1, FER3, FER4*, and *NEET* was not much induced in the *hy5* mutant unlike *pye* mutant. On the other hand, the *pye* mutant has high Fe content and the induction of Fe uptake related genes such as *FRO2, IRT1* was also not compromised in the *pye* mutant unlike *hy5* mutant which might be the reason for the enhanced expression of the Fe storage and assimilation related genes in the *pye* mutant (Figure S2). Only the Fe transport related genes such as *FRD3, YSL3, NPF5.9, OPT3, NAS4,* and *YSL2* were found to be commonly more upregulated in both the *pye* and *hy5* mutants as compared to the WT under Fe deficiency (Figure 3, S2). This suggests that HY5 might act together with PYE to balance the optimum induction of Fe transport under Fe deficiency. Overall, the phenotypic and transcriptomics data suggest that HY5 and PYE have some similar and some distinct roles in the regulation of Fe deficiency response.

Fe transport from root to shoot and its redistribution from shoots to various sink organs is very crucial for the supply of Fe to various plant parts and ultimately optimum plant growth. Fe is mainly transported and redistributed to different plant parts via xylem and phloem. To maintain Fe homeostasis under conditions of Fe deficiency, there is induction of Fe uptake, Fe transport, and repression of Fe storage. In our study, we found that key Fe transport genes including *FRD3, OPT3, NPF5.9, YSL3,* and *YSL2* were more induced in the *hy5* mutant as compared to the WT under -Fe conditions and HY5 directly binds to the promoter of these genes (Figure 3, 4). These results indicate that HY5 acts as a direct transcriptional repressor to regulate the optimum induction of Fe transport genes including *OPT3, FRD3, NPF5.9, YSL3,* and *YSL2* under -Fe. HY5 has been known to act as a transcriptional activator to positively regulate genes responsible for Fe uptake (*FRO2* directly and *IRT1, FIT* indirectly). Thus, despite the enhanced Fe transport in the *hy5* mutant, it is sensitive to Fe deficiency as the induction of Fe uptake is compromised.

PYE is known to interact with ILR3 and directly negatively regulate the expression of genes involved in Fe transport (*NAS4*), storage (*FER1, FER3, FER4* and *VTL2*) and assimilation *(NEET).* In addition to this, PYE is also known to repress its own expression (Tissot *et al*., 2019). In our study, we checked whether PYE also binds on the promoter of other Fe transport genes on the same region as HY5 does and we found that PYE does not bind to the promoter of Fe transport genes. In our previous study, we have shown that HY5 binds on the promoter of PYE and negatively regulate its expression (Mankotia *et al*., 2023). Here, we confirmed that PYE also binds to the same region as HY5 does on its promoter (Figure 6a). The *NAS4* gene expression was also found to be more induced in the *hy5* mutant as compared to the WT (Figure 3e). We found that HY5 also interacts on the same region of *NAS4* as PYE does and negatively regulates its expression (Figure 6b). These results suggest that some of the genes including *NAS4* and *PYE* are regulated by PYE and HY5 together, whereas the other Fe homeostasis related genes could be regulated by HY5 and PYE independently, possibly by interacting with separate activators and repressors involved in maintenance of Fe homeostasis.

Based on our results and existing literature, we propose a model (Figure 7) describing the role of HY5 in intercellular Fe transport under Fe deficiency in Arabidopsis. Importantly, we found that HY5 directly negatively regulates the expression of genes involved in Fe transport such as *OPT3, YSL2, YSL3, FRD3* and *NPF5.9*. Additionally, HY5 interacts with PYE and they both interact with the same region of *PYE* and *NAS4* promoter.

**Figure 7.**
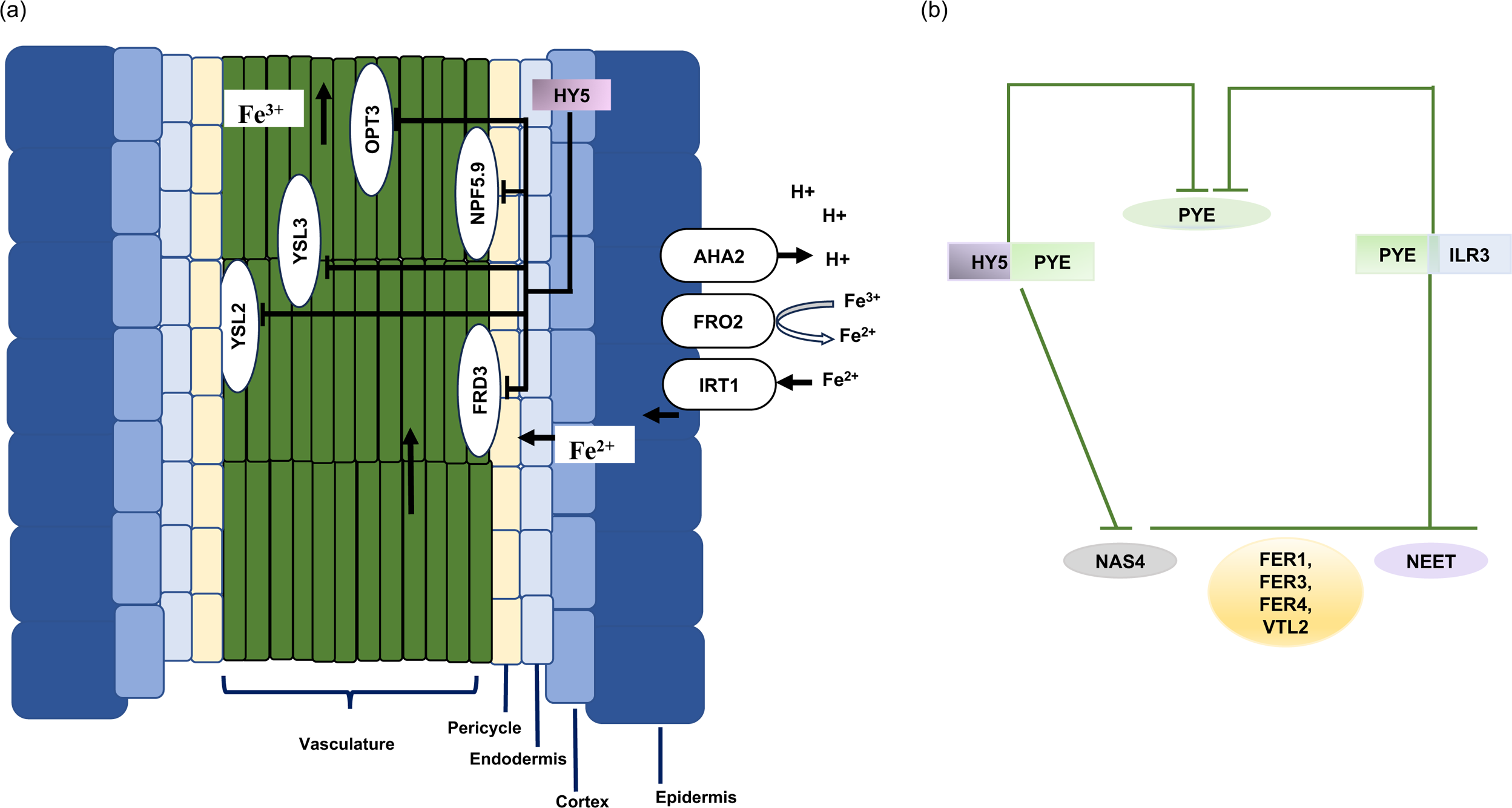
Model describing the role of HY5 in regulation of Fe homeostasis. **(a)** Under Fe deficiency, HY5 acts a direct transcriptional repressor of genes involved in intercellular transport of Fe from the vascular tissues such as *OPT3*, *FRD3*, *NPF5.9*, *YSL2* and *YSL3*. **(b)** HY5 interacts with PYE. PYE is known to interact with ILR3 and repress the expression of *NAS4* (involved in Fe transport), *FER1, FER3, FER4* and *VTL2* (involved in Fe storage) and *NEET* (involved in Fe assimilation) as well as it represses its own expression. PYE and HY5 interact at the same region on the promoter of *PYE* and *NAS4* to negatively regulate their expression. Oval boxes represent mRNA and rectangular boxes represent protein.

In summary, the data produced in this study and in the previous studies suggest HY5 interacts with PYE, and they play some common roles as well as additive or independent roles in regulation of Fe deficiency responses. HY5 and PYE were found to together regulate the expression of *NAS4* as well as *PYE*. HY5 acts as a both transcriptional activator and repressor of plant responses to Fe deficiency which ensures the optimum expression of genes involved in maintenance of Fe homeostasis which is crucial for plant growth and development. Further studies are required to find out what other genes are regulated by HY5 and PYE together and whether HY5 interacts with any other bHLH TFs to act as a transcriptional activator to regulate Fe deficiency response. The data presented in this study demonstrates that HY5 function is indispensable for seedling growth under conditions of Fe deficiency. HY5 orthologs are present in major food crops such as Rice, tomato and soybean and may play a similar role in Fe mobilization and transport. This study provides new insights into plant Fe nutrition and potential targets for improving Fe nutrition of plants, enhancing seed Fe contents and improving crop yield.

## ACKNOWLEDGMENTS

We thank Roman Ulm for providing *pHY5:HY5:YFP/hy5* (Ler) seeds. We are grateful to Terry A. Long for *pye*, and *pPYE:PYE:eGFP/pye* seeds. We are grateful to all the members of SBS laboratory, and Ajay Kumar Pandey for critical reading of the manuscript. S.B.S. acknowledges intramural funding support from Indian Institute of Science Education and Research (IISER) Mohali. S.B.S. is a recipient of Ramalingaswami Fellowship from Department of Biotechnology (DBT) Govt. of India and acknowledges funding received from DBT. SBS also acknowledges Science and Engineering Research Board (SERB) for research funding (CRG/223 2022/003773). The SBS laboratory is also supported by the Indo-French Centre for the Promotion of Advanced Research (IFCPAR/CEFIPRA) under project 68T06-1. SM and PJ acknowledge PhD fellowship received from IISER Mohali and CSIR-UGC respectively.

## CONFLICT OF INTEREST STATEMENT

The authors declare no conflict of interest.

## Notes

### Competing Interest Statement

The authors have declared no competing interest.

## REFERENCES

Ang, L., Chattopadhyay, S., Wei, N., Oyama, T., Okada, K., Batschauer, A. and Deng, X. (1998) Molecular Interaction between COP1 and HY5 Defines a Regulatory Switch for Light Control of Arabidopsis Development., 1, 213–222.

Aono, M., Kubo, A., Saji, H., Tanaka, K. and Kondo, N. (1993) Enhanced tolerance to photooxidative stress of transgenic nicotiana tabacum with high chloroplastic glutathione reductase activity. Plant Cell Physiol.

Balk, J. and Schaedler, T.A. (2014) Iron Cofactor Assembly in Plants. Annu. Rev. Plant Biol, 65, 125–153. Available at: www.annualreviews.org.

Bolger, A.M., Lohse, M. and Usadel, B. (2014) Trimmomatic: A flexible trimmer for Illumina sequence data. Bioinformatics, 30, 2114–2120.

Brumbarova, T., Bauer, P. and Ivanov, R. (2015) Molecular mechanisms governing Arabidopsis iron uptake. Trends Plant Sci., 20, 124–133. Available at: 10.1016/j.tplants.2014.11.004.

Burko, Y., Seluzicki, A., Zander, M., Pedmale, U. V., Ecker, J.R. and Chory, J. (2020) Chimeric activators and repressors define HY5 activity and reveal a light-regulated feedback mechanism. Plant Cell, 32, 967–983.

Chen, S.Y., Gu, T.Y., Qi, Z.A., Yan, J., Fang, Z.J., Lu, Y.T., Li, H. and Gong, J.M. (2021) Two NPF transporters mediate iron long-distance transport and homeostasis in Arabidopsis. Plant Commun., 2, 1–11.

Delker, C., Sonntag, L., James, G.V., et al. (2014) The DET1-COP1-HY5 Pathway Constitutes a Multipurpose Signaling Module Regulating Plant Photomorphogenesis and Thermomorphogenesis. Cell Rep., 9, 1983–1989.

DiDonato, R.J., Roberts, L.A., Sanderson, T., Eisley, R.B. and Walker, E.L. (2004) Arabidopsis Yellow Stripe-Like2 (YSL2): A metal-regulated gene encoding a plasma membrane transporter of nicotianamine-metal complexes. Plant J., 39, 403–414.

Durrett, T.P., Gassmann, W. and Rogers, E.E. (2007) The FRD3-mediated efflux of citrate into the root vasculature is necessary for efficient iron translocation. Plant Physiol., 144, 197–205.

Eide, D., Broderius, M., Fett, J. and Guerinot, M. Lou (1996) A novel iron-regulated metal transporter from plants identified by functional expression in yeast. Proc. Natl. Acad. Sci. U. S. A., 93, 5624–5628.

Fourcroy, P., Sisó-Terraza, P., Sudre, D., et al. (2014) Involvement of the ABCG37 transporter in secretion of scopoletin and derivatives by Arabidopsis roots in response to iron deficiency. New Phytol., 201, 155–167.

Gangappa, S.N. and Botto, J.F. (2016) The Multifaceted Roles of HY5 in Plant Growth and Development. Mol. Plant, 9, 1353–1365.

Gangappa, S.N. and Kumar, S.V. (2017) DET1 and HY5 Control PIF4-Mediated Thermosensory Elongation Growth through Distinct Mechanisms. Cell Rep., 18, 344–351. Available at: 10.1016/j.celrep.2016.12.046.

Gao, F., Robe, K. and Dubos, C. (2020) Further insights into the role of bHLH121 in the regulation of iron homeostasis in Arabidopsis thaliana. Plant Signal. Behav., 00. Available at: 10.1080/15592324.2020.1795582.

Gendrel, A.V., Lippman, Z., Martienssen, R. and Colot, V. (2005) Profiling histone modification patterns in plants using genomic tiling microarrays. Nat. Methods, 2, 213–218.

Grillet, L., Lan, P., Li, W., Mokkapati, G. and Schmidt, W. (2018) IRON MAN is a ubiquitous family of peptides that control iron transport in plants. Nat. Plants, 4, 953–963.

Hänsch, R. and Mendel, R.R. (2009) Physiological functions of mineral micronutrients (Cu, Zn, Mn, Fe, Ni, Mo, B, Cl). Curr. Opin. Plant Biol., 12, 259–266.

Haydon, M.J. and Cobbett, C.S. (2007) A novel major facilitator superfamily protein at the tonoplast influences zinc tolerance and accumulation in Arabidopsis. Plant Physiol., 143, 1705–1719.

Heazlewood, J.L., Tonti-Filippini, J.S., Gout, A.M., Day, D.A., Whelan, J. and Millar, A.H. (2004) Experimental Analysis of the Arabidopsis Mitochondrial Proteome Highlights Signaling and Regulatory Components, Provides Assessment of Targeting Prediction Programs, and Indicates Plant-Specific Mitochondrial Proteins. Plant Cell, 16, 241–256.

Kim, S.A., LaCroix, I.S., Gerber, S.A. and Guerinot, M. Lou (2019) The iron deficiency response in Arabidopsis thaliana requires the phosphorylated transcription factor URI. Proc. Natl. Acad. Sci. U. S. A., 116, 24933–24942.

Klatte, M., Schuler, M., Wirtz, M., Fink-Straube, C., Hell, R. and Bauer, P. (2009) The analysis of arabidopsis nicotianamine synthase mutants reveals functions for nicotianamine in seed iron loading and iron deficiency responses. Plant Physiol., 150, 257–271.

Kobayashi, T. and Nishizawa, N.K. (2012) Iron uptake, translocation, and regulation in higher plants. Annu. Rev. Plant Biol., 63, 131–152.

Li, X., Zhang, H., Ai, Q., Liang, G. and Yu, D. Two bHLH Transcription Factors, bHLH34 and bHLH104, Regulate Iron Homeostasis in Arabidopsis thaliana 1. Available at: www.plantphysiol.org/cgi/doi/10.1104/pp.15.01827.

Li, Y., Lu, C.K., Li, C.Y., et al. (2021) IRON MAN interacts with BRUTUS to maintain iron homeostasis in Arabidopsis. Proc. Natl. Acad. Sci., 118, e2109063118.

Liang, G., Zhang, H., Li, X., Ai, Q. and Yu, D. (2017) bHLH transcription factor bHLH115 regulates iron homeostasis in Arabidopsis thaliana. J. Exp. Bot., 68, 1743–1755. Available at: https://abrc.osu.edu/.

Liao, Y., Smyth, G.K. and Shi, W. (2014) FeatureCounts: An efficient general purpose program for assigning sequence reads to genomic features. Bioinformatics, 30, 923–930.

Long, T.A., Tsukagoshi, H., Busch, W., Lahner, B., Salt, D.E. and Benfey, P.N. (2010) The bHLH transcription factor POPEYE regulates response to iron deficiency in arabidopsis roots. Plant Cell, 22, 2219–2236.

Mankotia, S., Singh, D., Monika, K., Kalra, M., Meena, H., Meena, V., Yadav, R.K., Pandey, A.K. and Satbhai, S.B. (2023) ELONGATED HYPOCOTYL 5 regulates BRUTUS and affects iron acquisition and homeostasis in Arabidopsis thaliana. Plant J., 114, 1267–1284.

Masi, F. De, Grove, C.A., Vedenko, A., Alibés, A., Gisselbrecht, S.S., Serrano, L., Bulyk, M.L. and Walhout, A.J.M. (2011) Using a structural and logics systems approach to infer bHLH-DNA binding specificity determinants. Nucleic Acids Res., 39, 4553–4563.

Mukherjee, I., Campbell, N.H., Ash, J.S. and Connolly, E.L. (2006) Expression profiling of the Arabidopsis ferric chelate reductase (FRO) gene family reveals differential regulation by iron and copper. Planta, 223, 1178–1190.

Nawkar, G.M., Kang, C.H., Maibam, P., et al. (2017) HY5, a positive regulator of light signaling, negatively controls the unfolded protein response in Arabidopsis. Proc. Natl. Acad. Sci. U. S. A., 114, 2084–2089.

Nigel j. Robinson, Catherine M. Procter, Erin L. Connolly and Mary Lou Guerinot (1999) A ferric-chelate reductase for iron uptake from soils. Nature, 397, 694–697.

Petit, J.M., Briat, J.F. and Lobréaux, S. (2001) Structure and differential expression of the four members of the Arabidopsis thaliana ferritin gene family. Biochem. J., 359, 575–582.

Ravet, K., Touraine, B., Boucherez, J., Briat, J.F., Gaymard, F. and Cellier, F. (2009) Ferritins control interaction between iron homeostasis and oxidative stress in Arabidopsis. Plant J., 57, 400–412.

Ruckle, M.E., DeMarco, S.M. and Larkin, R.M. (2007) Plastid signals remodel light signaling networks and are essential for efficient chloroplast biogenesis in Arabidopsis. Plant Cell, 19, 3944–3960.

Santi, S. and Schmidt, W. (2009) Dissecting iron deficiency-induced proton extrusion in Arabidopsis roots. New Phytol., 183, 1072–1084.

Selote, D., Samira, R., Matthiadis, A., Gillikin, J.W. and Long, T.A. (2015) Iron-binding e3 ligase mediates iron response in plants by targeting basic helix-loop-helix transcription factors1[open]. Plant Physiol., 167, 273–286.

Stacey, M.G., Koh, S., Becker, J. and Stacey, G. (2002) AtOPT3, a member of the oligopeptide transporter family, is essential for embryo development in Arabidopsis. Plant Cell, 14, 2799–2811.

Stacey, M.G., Patel, A., McClain, W.E., Mathieu, M., Remley, M., Rogers, E.E., Gassmann, W., Blevins, D.G. and Stacey, G. (2008) The arabidopsis AtOPT3 protein functions in metal homeostasis and movement of iron to developing seeds. Plant Physiol., 146, 589–601.

Stark, C., Breitkreutz, B.J., Reguly, T., Boucher, L., Breitkreutz, A. and Tyers, M. (2006) BioGRID: a general repository for interaction datasets. Nucleic Acids Res., 34, 535–539.

Tissot, N., Robe, K., Gao, F., et al. (2019) Transcriptional integration of the responses to iron availability in Arabidopsis by the bHLH factor ILR3. New Phytol.

Vert, G., Grotz, N., Dédaldéchamp, F., Gaymard, F., Guerinot, M. Lou, Briat, J.F. and Curie, C. (2002) IRT1, an Arabidopsis transporter essential for iron uptake from the soil and for plant growth. Plant Cell.

Wang, N., Cui, Y., Liu, Y., Fan, H., Du, J., Huang, Z., Yuan, Y., Wu, H. and Ling, H.Q. (2013) Requirement and functional redundancy of Ib subgroup bHLH proteins for iron deficiency responses and uptake in arabidopsis thaliana. Mol. Plant, 6, 503–513. Available at: 10.1093/mp/sss089.

Waters, B.M., Chu, H.H., DiDonato, R.J., Roberts, L.A., Eisley, R.B., Lahner, B., Salt, D.E. and Walker, E.L. (2006) Mutations in Arabidopsis Yellow Stripe-Like1 and Yellow Stripe-Like3 reveal their roles in metal ion homeostasis and loading of metal ions in seeds. Plant Physiol., 141, 1446–1458.

Xu, D., Jiang, Y., Li, J., Lin, F., Holm, M. and Deng, X.W. (2016) BBX21, an Arabidopsis B-box protein, directly activates HY5 and is targeted by COP1 for 26S proteasome-mediated degradation. Proc. Natl. Acad. Sci. U. S. A., 113, 7655–7660.

Yadukrishnan, P., Rahul, P.V. and Datta, S. (2020) HY5 suppresses, rather than promotes, abscisic acid-mediated inhibition of postgermination seedling development. Plant Physiol., 184, 574–578.

Ying Yi and Mary Lou Guerinot (1996) The Plant Journal - November 1996 - Yi - Genetic evidence that induction of root Fe III chelate reductase activity is.pdf.

Yuan, Y., Wu, H., Wang, N., Li, J., Zhao, W., Du, J., Wang, D. and Ling, H.-Q. (2008) FIT interacts with AtbHLH38 and AtbHLH39 in regulating iron uptake gene expression for iron homeostasis in Arabidopsis. Cell Res., 18, 385–397. Available at: www.cell-research.com.

Zhang, L., Li, G., Wang, M., Di, D., Sun, L., Kronzucker, H.J. and Shi, W. (2018) Excess iron stress reduces root tip zone growth through nitric oxide-mediated repression of potassium homeostasis in Arabidopsis. New Phytol., 219, 259–274.

Zhang, X., Huai, J., Shang, F., Xu, G., Tang, W., Jing, Y. and Lin, R. (2017) A PIF1/PIF3-HY5-BBX23 transcription factor cascade affects photomorphogenesis. Plant Physiol., 174, 2487–2500.

Zhang, Y., Park, C., Bennett, C., Thornton, M. and Kim, D. (2021) Rapid and accurate alignment of nucleotide conversion sequencing reads with HISAT-3N. Genome Res., 31, 1290–1295.

