## Supplementary Figure 1, 2 for "ELONGATED HYPOCOTYL 5 (HY5) and POPEYE (PYE) Regulate Intercellular Iron Transport in Plants"

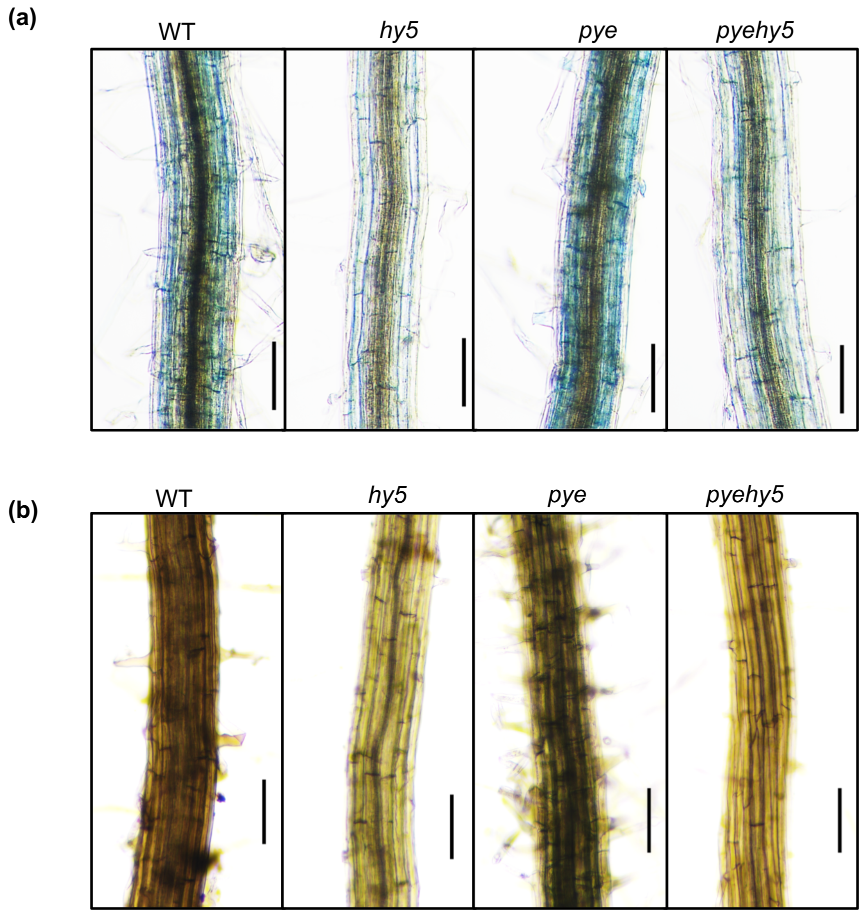


**Figure S1. (a)** Perl's stained and **(b)** Perls/Dab stained maturation zone of WT, *hy5, pye* and *pyehy5* grown on Fe-sufficient medium for 5 days. Bars = 100µm.

**(a)**

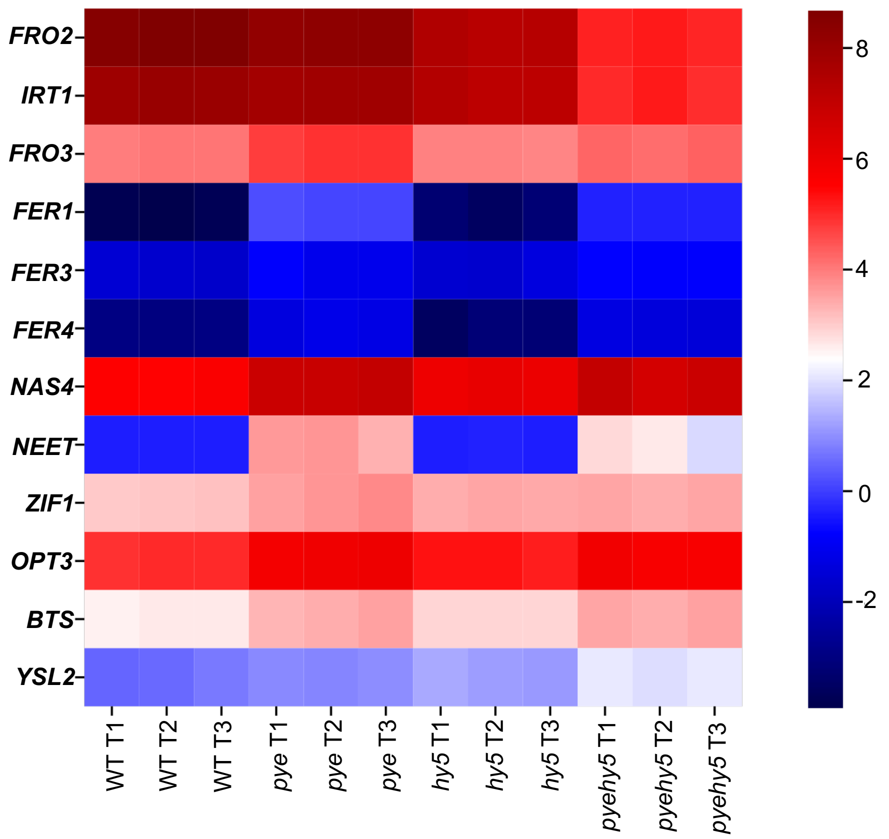


**Figure S2.** Heat map of selected Fe deficiency responsive genes. The color bar on the right side demonstrates the log2 (FoldChange) of a given gene in response to iron deficiency for a given genotype.
