## Supplemental Table 1 for "ELONGATED HYPOCOTYL 5 (HY5) and POPEYE (PYE) Regulate Intercellular Iron Transport in Plants"

Supplemental Table 1 Primers used in this study

| **Primer** | **Sequences** |
|---|---|
| *pye-1, LP* | TTCAAGACCTCATTCACTGGC |
| *pye-1, RP* | GGGGATTGATTATGTTTGGTG |
| *hy5, LP* | TTCACTCTCGATATCCGTTCG |
| *hy5, RP* | ATGCGAGTGAATGACCATTTC |
| qRT_*NAS4*_F | GGCTTCGACGTTGTGTTCTT |
| qRT_*NAS4*_R | AGCAAAGCACCAGGAGACAT |
| *qOPT3-F* | AAGCTTACTATAAACAGAGCCTTAGCTT |
| *qOPT3-R* | ACAGGATCAACAAGGTACCTCCTC |
| qRT *YSL2* FP | GGATACTTATTCTTCTCCCTTGTC |
| qRT *YSL2* RP | CCATCGTTTTTTCCTGCC |
| *pOPT3* Chip FP | CATACTCCTCTTAATAACATTGG |
| *pOPT3* Chip RP | CAGAAAGTGAATGCTGTTAC |
| *pOPT3* Chip II RP | CATTGGGAGGTTCCAAATGG |
| *pOPT3* Chip II FP | AGTGTCAAAAAAACGGGACC |
| *pOPT3* Chip III FP | TTATGTTCTGCGCACACCAC |
| *pOPT3* Chip III RP | TCCAGATCAAAGCTTGTCTC |
| *pOPT3* Chip IV FP | GGACAACCAATAGAAAGTGC |
| *pOPT3* Chip IV RP | CTAGGTGTGGTTAGCTCGTG |
| *pOPT3* Chip V FP | CAAGAGAGATTCATGCATGT |

| | |
|---|---|
| *pOPT3* Chip V RP | AACACAGTCTATGTTAGCTG |
| *pYSL2* ChIP FP | TTTTATGTTACCTCCTAACTTAC |
| *pYSL2* ChIP RP | CACTGTTACACAACAACAATATTT |
| *pFRD3* ChIP FP | GGAAACCTTTGTTTTCTTC |
| *pFRD3* ChIP RP | GAAGATATCAATAAGTGTTTCG |
| *pFRD3* ChIP II FP | GCGGTTAACTCTACGATAAC |
| *pFRD3* ChIP II RP | GTTGTATATAGTGCGTGTCG |
| *pYSL3* ChIP FP | TTCAGATAATTATGCTTGGG |
| *pYSL3* ChIP RP | TCAAAACAAAAACCCAAAAG |
| *pYSL3* ChIP II FP | ATGAAGGTTATATATGTGGAG |
| *pYSL3* ChIP II RP | AAGAGAAAAGAGTTTTGGGG |
| *pNPF5.9* ChIP FP | GTTTTTATATGCGACGCCTG |
| *pNPF5.9* ChIP RP | AGGTATCATGTACGAAGAGG |
| *pNPF5.9* ChIP II FP | AATCCACCCTATAATGGCAC |
| *pNPF5.9* ChIP II RP | CTTTCTTTTTTTGTCGAAAGG |
| *pPYE* ChIP FP | GACGTGTCCATGAGAGATGA |
| *pPYE* ChiP RP | TTTTTGGAGGAAGAAGGTCC |
| *pNAS4* ChiP FP | CGAAATATGAAGACAACACATGC |
| *pNAS4* ChiP RP | TGAGAGTACACGTGCCATCG |
